# Exploration of oncogenic cooperation between germline variation and somatic mutation in prostate cancer progression

**DOI:** 10.1101/2025.08.05.668679

**Authors:** Zuzanna A. Kołodyńska, Kimberley M. Hanssen, Florian H. Geyer, Maximillian M. L. Knott, Karim Aljakouch, Alina Ritter, Alexandra Delipetrou, Jeroen Krijgsveld, Erick Mitchell-Velasquez, Irfan Asangani, Rainer Will, Martina Müller-Nurasyid, Konstantin Strauch, Alexander Buchner, Florencia Cidre-Aranaz, Thomas G.P. Grünewald

## Abstract

Prostate carcinoma (PCa) is the most common cancer of men, associated with a still unresolved issue of accurate risk-stratification. While recent advances in omics technologies have provided clues as to how molecular changes shape the onset and progression of PCa, it remains largely unclear whether germline variants and somatic mutations cooperate to contribute to PCa progression and outcome. Thus, we explored whether oncogenic cooperation between regulatory germline variants and somatic driver mutations can help explain why some PCa patients develop a more aggressive phenotype, which may have implications for risk-adapted medical treatment. Here, by employing an integrative functional genomics approach, we identified *receptor-type protein-tyrosine phosphatase kappa* (*PTPRK*) as a TMPRSS2::ERG (TE)-modulated gene associated with PCa progression whose expression is controlled by cooperation of the TE-fusion with a regulatory single nucleotide polymorphism (SNP). Analysis of available clinically annotated patient cohorts demonstrated that PTPRK is overexpressed in TE-positive PCa tumors and associated with higher Gleason scores and metastatic disease. TE knockdown in PCa cell lines reduced PTPRK expression, while ectopic overexpression of the fusion in TE-negative PCa cell lines and prostatic epithelium cells induced its expression. Functionally, PTPRK silencing inhibited cellular proliferation, cell cycle progression, and clonogenic growth of PCa cells, which was mirrored by dysregulation of corresponding gene and protein signatures in global transcriptomic and phospho-proteomic analyses after PTPRK knockdown. Analysis of TE ChIP-Seq and Hi-C data from PCa cells highlighted a proximal TE-bound DNA element whose TE-dependent enhancer activity was validated in reporter assays and which could be abrogated by a regulatory SNP. Collectively, our results provide evidence of how exploration of oncogenic cooperation may help to identify novel biomarkers and potentially druggable pathways and highlight the role of the regulatory genome in PCa progression.

## LETTER

Prostate carcinoma (PCa) is the most common malignancy in men worldwide and presents with a highly variable clinical phenotype ranging from indolent to very aggressive disease (1). While active surveillance allows clinicians to monitor patients with early PCa without imposing unnecessary treatment and thus side-effects, this strategy bears the risk of overlooking aggressive tumors with potential for rapid disease progression (2). Hence, there is an urgent need to develop biomarkers to better distinguish tumors requiring immediate action from less aggressive forms that can enter active surveillance programs (3).

Genetically, ∼50% of PCa tumors are characterized by the *TMPRSS2::ERG* (TE) oncogene, formed by an intrachromosomal fusion of the androgen-regulated *TMPRSS2* gene and the ETS transcription factor *ERG* located on chr21 (4). Biologically, TE is associated with enhanced invasion and increased cell proliferation of PCa cells, thus conferring a more aggressive phenotype to preclinical models (5)(6). Although TE-positive and -negative PCa are clearly distinct molecular subtypes with different epigenetic DNA-methylation profiles (7)(8), 20 years after its discovery (4) the clinical/prognostic value of the fusion remains controversial. While some studies report that patients with TE-positive tumors have reduced survival compared to TE-negative tumors (9)(10)(11), other studies have found that TE expression may not be associated with biochemical recurrence (12) or patient outcome (13) and even be linked with a more favorable prognosis (14)(15). These contrasting results indicate that although the TE fusion has a functional impact on PCa biology, its presence alone may be insufficient to constitute a useful biomarker for assessment of clinical PCa aggressiveness. In support of this notion, a recent study demonstrated that the prognostic value of other molecular PCa biomarkers strongly depends on the presence/absence of TE (16), suggesting that the fusion may interact or cooperate with additional factors that need to be considered for accurate interpretation of its biological and/or clinical relevance. Recent evidence suggest that the evolution of the somatic mutational landscape in PCa is shaped by the germline context (17). Likewise, an important earlier study highlighted that the set of PCa germline risk variants identified by genome-wide associations studies (GWAS) were distinct in PCa patients whose tumors harbored the TE fusion versus those that lacked TE (18). Similarly, another recent study demonstrated that germline variants may modulate the transcriptional output of TE in PCa (19). However, while several GWAS have identified dozens of germline loci which may contribute to PCa susceptibility and/or aggressiveness (20)(21)(22)(23), most of these studies were TE-agnostic.

In contrast, in Ewing sarcoma – a pediatric bone or soft-tissue cancer driven by EWSR1::FLI1 or EWSR1::ERG fusions (24) – accumulating evidence demonstrates an important role of oncogenic cooperation of regulatory germline variants with the respective fusions that governs expression of genes involved in Ewing sarcoma susceptibility and/or aggressiveness (25)(26)(27). Thus, to investigate if such oncogenic cooperation may also take place in PCa and whether its exploration can yield additional biomarkers and/or targets, we employed an integrative genetic and functional approach. To identify clinically relevant TE target genes whose expression may be the subject of oncogenic cooperation between regulatory (germline) variants and the TE fusion, we crossed multiple publicly available datasets **(Fig. 1a)**. Firstly, we assessed transcriptome profiling data of the VCaP PCa cell line with/without conditional shRNA-mediated knockdown of TE (GSE60771)(6). Data normalization yielded a total of 19,702 genes, of which 4,669 were significantly differentially expressed upon TE silencing (|log_2_ FC| >0.5; *P≤*0.05; 4,417 downregulated and 252 upregulated by TE).

**Figure 1.**
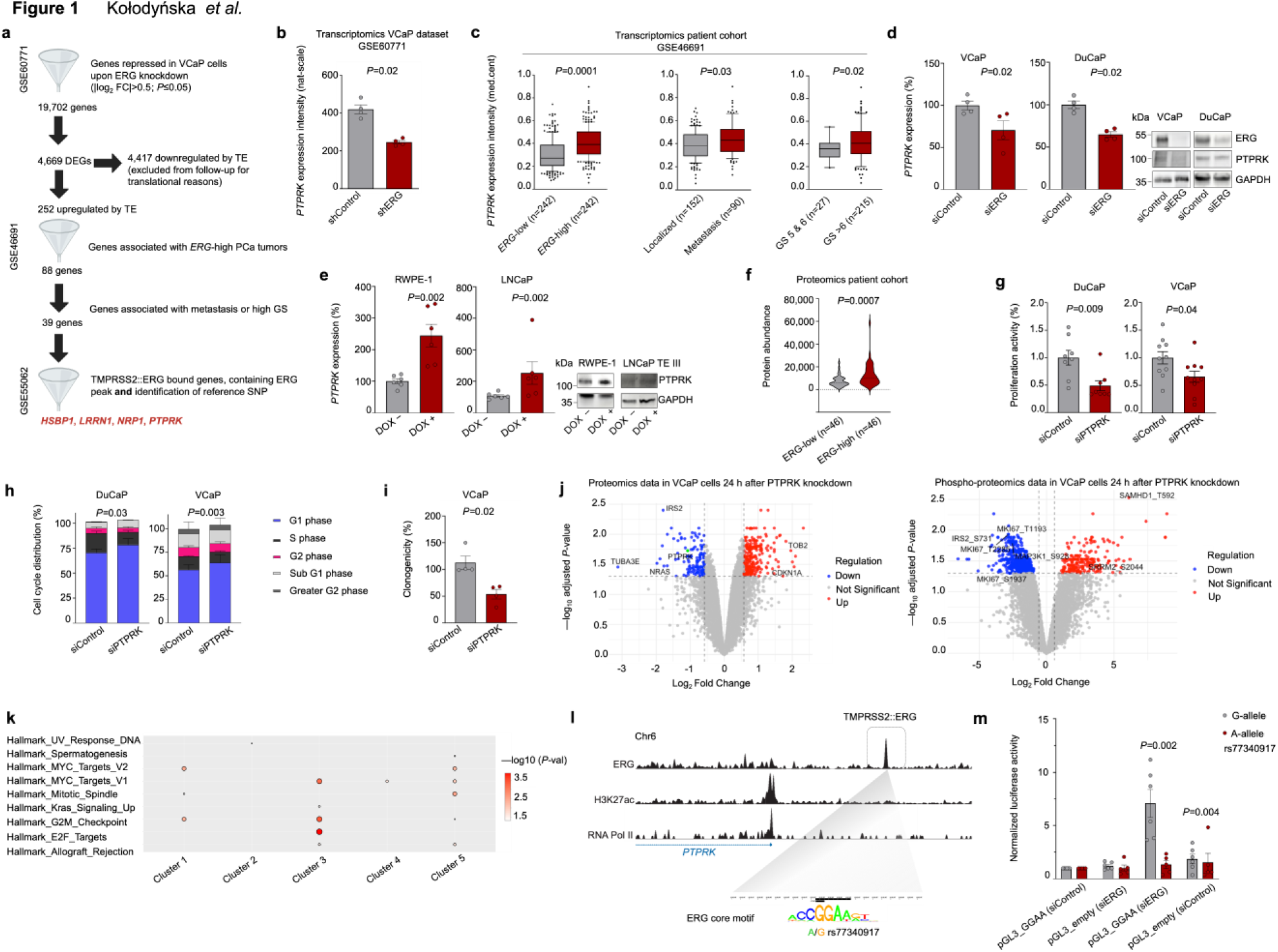
PTPRK is a TE-modulated gene associated with PCa progression, whose expression is controlled by cooperation of TE-fusion with a regulatory SNP. **a.** Workflow diagram of the pipeline cross-linking several public datasets to identify clinically relevant genes driven by TE and associated with metastasis and high GS in PCa (|log_2_ FC|>0.5; *P*≤0.05) **b.** Bar plot illustrating decreased expression of *PTPRK* following TE (*ERG*) silencing in VCaP cells, n=4 biologically independent experiments. Horizontal bars represent mean, whiskers represent the SEM. Two-sided Mann-Whitney test. **c.** Analysis of *PTPRK* expression in PCa patient tumors stratified by the indicated parameter. Two-sided Mann-Whitney test. **d.** Bar plot illustrating relative mRNA expression of *PTPRK*, quantified by qRT-PCR in VCaP and DuCaP PCa cell lines 72 h after transfection with siERG, n=4 biologically independent experiments. Horizontal bars represent mean, whiskers represent the SEM. ERG and PTPRK knockdown were confirmed *via* western blot of 20 μg whole cell lysate (loading control: GAPDH). Two-sided Mann-Whitney test. **e.** Bar plot illustrating relative mRNA expression of *PTPRK*, quantified by qRT-PCR in RWPE-1 and LNCaP cells 72 h after induction of *ERG* expression (DOX), n=6 biologically independent experiments. Horizontal bars represent mean, whiskers represent the SEM. ERG and PTPRK knockdown were confirmed *via* western blot of 20 μg whole cell lysate (loading control: GAPDH). Two-sided Mann-Whitney test. **f.** Violin plot illustrating PTPRK protein expression in PCa patient samples. The dotted line represents the median. Two-sided Mann-Whitney test. **g.** Analysis of cell proliferation of DuCaP and VCaP PCa cells 72 h after knockdown of *PTPRK*. Horizontal bars represent mean, whiskers indicate SEM, n=8−10 biologically independent experiments. Two-sided Mann-Whitney test. **h.** Analysis of the cell cycle in DuCaP and VCaP cells determined by PI staining and flow cytometry 72 h after *PTPRK* silencing. Horizontal bars represent mean, whiskers indicate SEM, n=5−6 biologically independent experiments. Unpaired two-sided student’s t-test. **i.** Colony-forming assay of VCaP cells, 7 d following transfection with siRNA targeting *PTPRK*. Horizontal bars represent mean, whiskers indicate SEM, n=4 biologically independent experiments. Two-sided Mann-Whitney test. **j.** Volcano plot (log_2_ protein fold change *vs P-* adjusted value) showing differentially expressed proteins in the whole proteome in response to PTPRK knockdown in VCaP cells after 24 h. Highlighted are proliferation and cell cycle-related proteins, including PTPRK highlighted in green (left). Volcano plot of the phosphoproteomic data (log_2_ protein fold change *vs P-*adjusted value). Phosphosites whose corresponding proteins are regulated at the proteome level are marked in blue (downregulated), red (upregulated) or grey (not significant). Highlighted are proliferation related proteins (right). **k.** Cancer Hallmark pathways from clustering of time series gene expression patterns (cluster 1−5) affected by PTPRK knockdown after 24 h in VCaP cells. **l.** Integrative genomic view of the *PTPRK* locus showing evidence of binding of TE close to promoter regions (hg19, chr6:128,861,959−128,862,733) from VCaP cell line. SNP G>A rs77340917 is located central to TE-peak which disturbs the ERG-binding motif. ChIP-seq signals (GSE55062) are depicted for TE, H3K27ac and RNA polymerase II. **m.** Luciferase reporter assay signal of the pGL3 vector containing the G or A allele at rs77340917 SNP. Horizontal bars represent mean, whiskers indicate SEM, n=5−6 biologically independent experiments. Two-sided Mann-Whitney test with Bonferroni correction for multiple comparisons.

Since clinical targeting of candidate genes may be more feasible if they are overexpressed in the respective cancer, we subsequently focused on the 252 TE-induced differentially expressed genes (DEGs). These 252 genes were checked for two additional aspects: 1) whether they are overexpressed in TE-positive versus -negative patient tumors, and 2) whether their overexpression is linked with clinicopathological parameters associated with more aggressive disease. To that end, we utilized a large gene expression dataset from 639 PCa patients that contained information on the disease state (localized versus metastatic) and Gleason score (GS) as surrogate markers for biological behavior of the tumors (GSE46691) (28). Since this dataset did not contain information on somatic mutations and previous studies had demonstrated an excellent correlation of the fusion status with *ERG* expression levels (29)(30), we inferred the TE status from *ERG* expression levels (see Methods). This process yielded 484 PCa patients with clinicopathological annotations and gene expression data stratified into 242 *ERG*-high and 242 *ERG*-low cases. Crossing the 252 downregulated genes upon TE knockdown in the VCaP model with these patient data showed that of those 252 genes, 88 exhibited a significantly higher expression in *ERG*-high compared to *ERG*-low PCa tumors. Next, we tested whether the expression of those 88 genes were different between patients that presented with localized or metastatic disease at diagnosis or with high GS (>6). Altogether this yielded a set of 39 genes that appeared to be regulated by TE in the VCaP model as well as in patient tumors and which were associated with relevant outcome parameters **(Supplementary Table 1)**. Finally, to identify direct TE targets that may be subject to modulation by a regulatory variant, we used publicly available ChIP-Seq data of the TE fusion to scan for signals mapping to proximal enhancers that may be modulated by common SNPs located close to or even within the ETS-like core motif for TE binding (GSE55062)(31). Inspection of the data in the UCSC genome browser highlighted four genes (*HSBP1*, *LRRN1*, *NRP1*, *PTPRK*) as promising candidates for functional follow-up. An example of the discovery analysis of one of these genes – *receptor-type protein-tyrosine phosphatase kappa* (*PTPRK*) – is outlined in **Figs. 1b, c.** To validate the *in silico* results and to test the biological relevance of these genes in PCa, we employed proliferation assays in VCaP PCa cells and the prostatic intraepithelial neoplasia cell line RWPE-1 **(Supplementary Figs. 1–3)**. Only *PTPRK* exhibited a consistent pattern of regulation by TE in functional validation experiments using the TE-positive VCaP and its derivative DuCaP cell lines with/without siPOOL-mediated knockdown of *ERG* **(Fig. 1d)** as well as in conditional heterologous ERG-overexpression models, including RWPE-1 and the fusion-negative PCa cell line LNCaP **(Fig. 1e)**. Thus, we henceforth focused on PTPRK, which has been previously reported to be overexpressed in PCa (32).

In line with our functional ERG-silencing and -overexpression experiments, reanalysis of a recently published proteomics dataset of PCa patients (33) showed that PTPRK is highly expressed at the protein level and significantly higher expressed in ERG-high versus -low PCa tumors **(Fig. 1f)**. RNAi-mediated silencing of *PTPRK* in VCaP and DuCaP cells demonstrated that this protein promotes cell proliferation, cell cycle progression and clonogenic growth of PCa cells **(Figs. 1g–i)**. These phenotypes were in agreement with data from combined transcriptomic, global proteomic and phospho-proteomic profiling of VCaP cells with/without siPOOL-mediated PTPRK knockdown after 24 h, which mainly showed a dysregulation of gene signatures and/or proteins, such as IRS2, NRAS, CDKN1A and Ki-67 involved in cell cycle regulation and proliferation **(Figs. 1j, k, Supplementary Table S2, S3, S4, Supplementary Figs. 4, 5)**. Thus, PTPRK appeared as a clinically relevant TE-driven gene affecting multiple downstream signalling pathways that converge in a more aggressive phenotype.

As shown in **Fig. 1l**, TE binds to a DNA element around ∼20,000 bp upstream of the transcriptional start site of *PTPRK* in VCaP cells. Underlying this ERG ChIP-Seq signal, we identified a consensus ETS binding site with the canonical GGAA core motif that contains a potential regulatory SNP G>A rs77340917 (**Fig. 1l**). Analysis of available Hi-C data from PCa cell lines (GSE172097) (34) confirmed that this DNA element physically interacts with the *PTPRK* promoter region **(Supplementary Fig. 6)**. To assess the potential TE-dependent enhancer activity of this DNA element and the potential regulatory effect of this SNP, we cloned this DNA element from VCaP cells, used site-specific mutagenesis to introduce this SNP, and carried out dual luciferase reporter assays in VCaP cells with/without siPOOL-mediated knockdown of TE (ERG). Strikingly, this DNA-element showed strong TE-dependent enhancer activity, which could be completely abrogated either by TE silencing or introduction of the SNP disrupting the GGAA core motif **(Fig. 1m)**. Such effect was not observed at another more distal TE-signal, which showed no physical interaction with the PTPRK promoter in the Hi-C data **(Supplementary Fig. 7)**.

Collectively, these results suggested that *PTPRK* is a TE-regulated gene related to PCa progression, whose expression levels may be regulated by at least one proximal ETS-like enhancer element that can be subject to germline variation. Thus, these data further establish the relevance of oncogenic cooperation between regulatory germline SNPs and somatic mutations (here TE) for tumor progression and heterogeneity in PCa patients and suggest that PTPRK may serve as a prognostic biomarker in PCa.

## Supporting information

Kolodynska_et_al_Supplementary Files

## AUTHOR CONTRIBUTIONS

Z.K., K.M.H. F.H.G, M.M.L.K. and A.R. performed experimental assays. K.A., A.D. and J. K. caried out proteomics analyses. E.M.-V. and I.F. provided Hi-C and ChIP-Seq data. R.W. carried out lentiviral transductions of cell line models. M.M.-N., K.S., A.B. and F.C.A. provided experimental and statistical guidance and/or were involved in funding acquisition and project discussion. Z.K. and T.G.P.G. wrote the manuscript and designed the figures and tables. T.G.P.G. conceived and supervised the study and provided the laboratory infrastructure and financial support. All authors read and approved the final manuscript.

## ACKNOWLEDGEMENTS

We thank Stefanie Kutschmann, Felina Zahnow, and Nadine Gmelin (Division of Translational Pediatric Sarcoma Research, DKFZ, Heidelberg) for excellent technical assistance. We thank the DKFZ NGS core facility for excellent service, and Prof. Dr. Holger Sültmann (Division of Cancer Genome Research, DKFZ, Heidelberg) for providing the DOX-inducible ERG overexpressing LNCaP cells. We thank Dr. Hayley J. Sharpe (Babraham Institute, Cambridge, United Kingdom) for kindly providing the PTPRK antibody as well Prof. Isabel Heidegger-Pircher (University of Innsbruck, Austria) for providing DuCaP cells.

## MATERIALS AND METHDOS

### Provenience of cell lines and cell culture conditions

Human cell lines were acquired from the following suppliers: RWPE-1 (RRID:CVCL_3791), VCaP (RRID:CVCL_2235), and HEK293-FT cells (RRID:CVCL_6911) were from the American Type Culture Collection (ATCC, Manassas, VA, USA); DuCaP (RRID:CVCL_2025) were kindly provided by Prof. Isabel Heidegger-Pircher (University of Innsbruck, Austria); LNCaP expressing TE III variant were kindly provided by Prof. Dr. Holger Sültmann (German Cancer Research Center, Heidelberg, Germany) (35).

For all experiments, DuCaP and LNCaP were grown in RPMI 1640 medium (Gibco) supplemented with 10% fetal bovine serum (FBS), 100 U/mL penicillin and 100 μg/mL streptomycin (Sigma-Aldrich), and 1% GlutaMAX (Gibco). VCaP and HEK293-FT cells were cultured in DMEM medium (Gibco) supplemented with 10% FBS and 100 U/mL penicillin and 100 μg/mL streptomycin. All cells were grown at 37 °C in a humidified 5% CO_2_ incubator. Cells were routinely checked for the absence of Mycoplasma contamination (Mycoplasma PCR Detection Kit, Applied Biological Materials) and cell line identity was verified by Short Tandem Repeat (STR) and/or single nucleotide polymorphism (SNP) profiling (Multiplexion, Heidelberg, Germany).

### Extraction of RNA, reverse transcription and quantitative real-time PCR (qRT-PCR)

Total RNA was isolated from cells using the NucleoSpin RNA kit (Macherey-Nagel GmbH & Co. KG Düren, Germany) according to the manufacturer’s instructions. RNA concentration was measured using NanoDrop One/One UV-Vis spectrophotometer (Thermo Fisher Scientific). RNA (1 μg) was utilized for complementary DNA (cDNA) synthesis using the High-Capacity cDNA Reverse Transcription Kit (Applied Biosystems, Waltham, MA, USA). qRT-PCR reactions were performed using the SYBR Select Master Mix for CFX (Applied Biosystems) in a final volume of 15 μL. All primer sequences used for qRT-PCR are listed in **Supplementary Table S5**. For each reaction, gene expression was normalized to *RPLP0* as an internal control using the ΔΔCt method. The qRT-PCR was performed using the BioRad CFX Connect instrument (Bio-Rad Laboratories GmbH, Feldkirchen, Germany) under the following conditions: UDG-Activation at 50 °C for 2 min, 95 °C for 2 min, followed by DNA denaturation at 95 °C for 15 sec with annealing and elongation at 60 °C for 1 min for a total of 40 cycles.

### RNA interference-mediated gene knockdown

PCa cells were seeded in 6-well plates at 1–8×10^5^ cells per well. Cells were transfected with siPOOLs (siTOOLs, Planegg, Germany) targeting *ERG* or *PTPRK* at a concentration of 10 nM for 72 h using Lipofectamine RNAiMAX (Thermo Fisher Scientific) according to manufacturer’s instructions. Each siPOOL consisted of 30 predefined specific siRNAs against the respective target, which maximizes on-target while minimizing potential off-target effects (36).

### Western blot

For each sample, a total of 20 μg of protein was prepared for the western blot. Whole cell lysate was extracted using RIPA buffer (SERVA Electrophoresis GmbH, Heidelberg, Germany) mixed with 1× Halt^TM^ Protease and Phosphatase Inhibitor Cocktail (Thermo Scientific). Analysis of protein concentration was determined using the Pierce BCA Protein Assay Kit (Thermo Scientific). The samples were separated on a 10% Tris-HCl gels at 120 V and transferred onto a PVDF membrane using the Trans-Blot Turbo Transfer System (BioRad). Following the transfer, the membranes were blocked in 5% milk in 1× TBS-T at room temperature for 1 h and incubated overnight at 4 °C with recombinant monoclonal anti-human PTPRK primary antibody (1:2,000) established and kindly provided by the Hayley J. Sharpe laboratory (Babraham Institute, Cambridge, United Kingdom) (37), mouse monoclonal anti-ERG (1:3,000, ab92513, abcam) or rabbit monoclonal anti-GAPDH (1:1,000, #2118, Cell Signaling Technology). Thereafter, the membranes were washed twice in TBS-T for 10 min each and incubated for 1 h with horseradish peroxidase (HRP) coupled secondary antibody mouse anti-rabbit IgG light chain (1:2,000, abcam, SB62a) or donkey anti-human IgG heavy and light chain (1:3,000, 709-035-149, Jackson ImmunoResearch Labs). Membranes were developed using chemiluminescence and Immobilon Western HRP Substrate (Sigma-Aldrich) and visualized using the FusionFX western blot & Chemi imaging system (Vilber).

### Generation of conditional *ERG* overexpression in RWPE-1 cell line

To conditionally (over)express *ERG*, we transduced the prostatic epithelial RWPE-1 cell line with lentivirus containing a doxycycline (DOX)-inducible *ERG* transcript variant (RefSeq NM_182918.4) 1 cloned into the pLIX_403 vector (Addgene, #41395) in collaboration with the DKFZ Cellular Tools Core Facility. To that end, the *ERG* transcript variant 1 was cloned into a Gateway entry vector (pDONR223) carrying a spectinomycin selection marker. The insert was confirmed with Sanger Sequencing (see **Supplementary Table S5** for the sequencing primer list). Lentiviral particles were synthesized using the standard protocol. In brief, HEK293-FT cells were transfected with packaging plasmids VSV.G (Addgene, plasmid #41395, Cambridge, MA, USA) and psPAX2 (Addgene, plasmid #12260, Cambridge, MA, USA). After 48 h, the viral supernatant was collected and filtered through a 0.45 μm filter. RWPE-1 cells were transduced for 24 h in a 6-well plate. After viral removal, transduced cells were selected with puromycin (1.5 μg/mL, InvivoGen). Overexpression of *ERG* was induced by adding 1 μg/mL of DOX to the medium and confirmed by qRT-PCR and immunoblotting.

### Analysis of publicly available datasets

Several publicly available datasets were downloaded and utilized from the Gene Expression Omnibus (GEO; accession codes: GSE60771, GSE46691, GSE55062). To investigate the potential regulatory relationship of *TMPRSS2::ERG* and *PTPRK*, we used microarray data (GSE60771, Affymetrix Human Genome U133 Plus 2.0 Array) of VCaP PCa cells with conditional DOX-inducible shRNA against *ERG* (6). These datasets were normalized using robust multi-array average (RMA) (38) and hybrid brainarray chip-description files (CDF) yielding one optimized probe-set per gene (39).

To correlate the levels of *PTPRK* expression with clinicopathological parameters, such as the presence of metastases at diagnosis or GS, we took advantage of a comprehensive dataset of 639 patients who underwent radical prostatectomy (GSE46691; Affymetrix Human Exon 1.0 ST Array) (28). This dataset was normalized using the SCAN algorithm (40) with brainarray CDF. Since this dataset did not contain information on the precise *TMPRSS2::ERG* fusion status, we inferred the fusion status from the detected *ERG* expression levels as previous studies have demonstrated an excellent correlation of the fusion status with *ERG* expression levels (29)(30). As described previously (16), patients were classified as either *ERG*-high or *ERG*-low if their intratumorally *ERG* expression levels ranged in the lower 45% or upper 45% of *ERG* expressors, respectively. Accordingly, the middle 10% of *ERG* expressors were intentionally excluded to avoid any misclassification of cases with intermediate *ERG* expression levels. To qualify genes as direct targets of TE, ChIP-seq data for TE enrichment was utilized (GSE55062) (31). The ChIP-seq signal intensities were displayed in UCSC Genome Browser and manually inspected for TE-signals at potential regulatory DNA elements. To examine the physical interaction of TE with the *PTPRK* promotor, we used publicly available Hi-C data (GSE172097)(34). To visualize combined ChIP-seq and Hi-C chromatin interaction, UCSC Genome Browser was used.

### Assessment of cell proliferation

To study the potential effect of PTPRK on cellular proliferation, PCa cells were seeded in triplicate wells of 6-well plates (DuCaP, VCaP: 8×10^5^ cells per well) and transfected with siPOOLs targeting *PTPRK* or siControl for 72 h using Lipofectamine RNAiMAX (Invitrogen). At 72 h post transfection, cells including their supernatants were collected and manually counted in standardized hemocytometers (Neubauer Improved, Hartenstein) using the Trypan Blue exclusion method (1:1 ratio). The number of viable (Trypan Blue-negative) and dead cells (Trypan Blue-positive) was recorded separately.

### Analysis of cell cycle

To investigate whether PTPRK silencing affects cell cycle, DuCaP and VCaP (8×10^5^ cells per well) were seeded in 6-well plates and transfected with 10 nM siPOOLs targeting *PTPRK* or siControl. At 72 h post-transfection, cells were fixed in 70% ice-cold ethanol and incubated for 45 min with 100 μg/mL RNAse (Thermo Fisher) and 0.5 mg/mL PI solution (BioLegend). Then, samples were assayed with BD FACS Canto Flow Cytometer (BD Sciences) and the analysis was performed using FlowJo v.10.10 software. An example of cell population gating is shown in **Supplementary Fig. 8**.

### Colony formation assay (CFA)

For CFAs, VCaP cells were seeded in duplicate wells (5×10^4^ cells per well) in 12-well plates and grown in their standard medium for 7 days. PCa cells were treated with siPOOLs targeting *PTPRK* or siControl (10 nM; re-transfection every 72 h). Seven days after transfection, the colonies were fixed with ice-cold methanol for 10 min and stained with crystal violet (CV) (Sigma-Aldrich) for another 10 min. Thereafter, the colonies were washed with distilled deionized water and left to dry. To quantify the clonogenic cell growth, the percentage of area covered by CV-stained colonies was multiplied by the count of colonies using Fiji/Image J software (41).

### RNA-sequencing (RNA-seq) and differential gene expression analyses

To assess the impact of *PTPRK* on gene expression in PCa cells, RNA-seq analysis was performed. To this end, VCaP and DuCaP PCa cell lines were seeded in 6-well plates and transfected with either siRNA targeting *PTPRK* or siControl for 72 h. Thereafter, total RNA was extracted with NucleoSpin RNA kit (Macherey-Nagel) according to the manufacturer’s instructions. RNA concentration was measured using NanoDrop One/One UV-Vis spectrophotometer (Thermo Fisher Scientific). Sequencing of the samples was performed at the DKFZ NGS Core Facility (Heidelberg, Germany) using Illumina NovaSeq 6000 Sequencing System (100 bp paired-end sequencing). To identify DEGs, standard workflow of R *DeSeq2* package version 1.40.2 was utilized.

### Fast gene-set enrichment analyses (fGSEA)

To identify biological pathways that are significantly enriched in a ranked list of genes, fast pre-ranked gene-set enrichment analysis (fGSEA) was performed in R using *fgsea* package version 1.26.0. fGSEA was completed on the hallmark gene set (H) and pathways from the KEGG database from the Human Molecular Signatures Database (MSigDB, v2023.2.Hs). The enriched gene sets were filtered for significance (adjusted *P*<0.01, |normalized enrichment score| >1.0) and plotted in R.

### Automated sample preparation (autoSP3) for proteome profiling

In brief, VCaP (5×10^8^ cells per well) cells were grown in 10 cm cell culture dishes and transfected with siPOOLs targeting *PTPRK* or siPOOLs targeting control for 24 h. Cell pellets were lysed in 75 μL RIPA lysis buffer (Thermo Fisher) followed by protein extraction using the AFA-ultrasonication using LE220R-plus ultrasonicator (Covaris Ltd, UK). Protein concentration was determined using BCA Protein Assay Kit according to the manufacturer’s instructions. For protein extraction and purification 100 μg of protein per samples was directly processed using the automated SP3 (autoSP3) protocol. This protocol, including protein clean-up, reduction and alkylation (using 10 mmol/L TCEP and 40 mol/L CAA at final concentration), and digestion (trypsin/protein ratio of 1:20) in 50 mM TEAB was performed on the Bravo liquid handling system (Agilent Technologies) (42). A fraction of the peptide mixture was used for global proteome profiling while the remaining was further dried by vacuum centrifugation at 45 °C.

### Automated Fe (III)-IMAC-based workflow for phosphopeptide enrichment

For phosphoproteomics, phosphorylated peptides were enriched using the Fe (III)-NTA (5 μL) cartridges (Agilent technologies) in an automated fashion using the AssayMAP Bravo Platform (Agilent Technologies) according to the Phosphopeptide enrichment V2.1 protocol provided by Agilent. Briefly, peptides were reconstituted in 80% ACN + 0.1% TFA in H_2_O and vortexed for 5 min. The cartridges were primed with 100 μL of 50% ACN + 0.1% TFA in H_2_O priming buffer (flow rate of 300 μL/min) and equilibrated with 50 μL of 80% ACN + 0.1% TFA in H_2_O equilibration buffer (flow rate of 10 μL/min). Next, peptides were loaded onto the cartridge at 3 μL/min flow rate. The cup wash, internal cartridge wash and stringent syringe wash steps were performed according to the protocol’s default settings. Phosphorylated peptides were eluted from the cartridges using 1% aqueous ammonia + 5% ACN in H_2_O at flow rate of 5 μL/min. Samples were dried by vacuum centrifugation at 45 °C, reconstituted in 0.1% FA, and analyzed by LC-MS/MS.

### Proteomic data acquisition

Global proteome and phosphoproteome samples were analyzed on a timsTOF Pro (Bruker Daltonics) coupled with an Easy-nLC 1200 system (Thermo Scientific). Peptides were separated over an 80-min gradient at 300 nL/min using solvent A (0.1% formic acid in ULC-MS grade water) and solvent B (0.1% formic acid in 80% acetonitrile in ULC-MS grade water). Data were acquired in DIA-PASEF mode, with full MS scans spanning 100–1,700 m/z and a 1/k₀ range of 0.65–1.42 Vs/cm² using a 100 ms ramp time. The duty cycle was fixed at 100%, ion polarity was set to positive, and TIMS mode was enabled. Collision energy was set to 1/k0 range from 0.65 to 1.42 Vs/cm^2^. For the DIA scans, a custom isolation window pattern was optimized, covering the precursor range of 377−1,194 m/z, mobility range 1/k0 range from 0.67 to 1.39 Vs/cm^2^, and cycle time estimate of 1.58 s.

### Data processing

Global proteome raw MS data were analyzed using DIA-NN 2.2.0 in library-free mode using the default settings with the Homo sapiens (Taxon ID: 9606) reviewed Swiss-Prot database. In brief, ASTA-based library-free digestion, as well as deep learning-based prediction of retention times and ion mobilities, was enabled. Carbamidomethylation of cysteines was set as a fixed modification, allowing a maximum of one variable modification. Trypsin/P was selected as the protease, permitting up to two missed cleavages. The match-between-runs (MBR) function was enabled. Phosphoproteome raw MS data were analyzed using Spectronaut 19.1 (Biognosys) with the library-free DirectDIA+ (Deep) workflow and Homo sapiens (Taxon ID: 9606) reviewed Swiss-Prot database. The default BGS Phospho PTM Workflow settings were applied. Acetyl (Protein N-term), Oxidation (M), Phospho (STY) were selected as variable modifications. Carbamidomethyl (C) was selected as fixed modifications. Trypsin/P was selected as the digestion enzyme, allowing up to two missed cleavages. Peptides with lengths ranging from a minimum of 7 to a maximum of 52 amino acids were considered. Site localization probability filter was applied to retain only phosphorylation sites with a localization probability greater than 0.75. Raw data were analyzed using the SmartPhos R package for joint analysis of proteomics and phosphoproteomics data, available under https://github.com/Lu-Group-UKHD/SmartPhos.

### Luciferase reporter assay

For luciferase reporter assays, a 775 bp fragment (hg19, chr6:128,861,959−128,862,733) encompassing the *PTPRK*-associated potential regulatory element was cloned from the VCaP PCa cell line using Platinum SuperFi II Green PCR Mastermix (Thermo Fisher). Primer sequences are listed in **Supplementary Table S5**. The PCR protocol included the initial denaturation for 30 sec at 98 °C; denaturation at 98 °C for 10 sec; annealing at 60 °C for 10 sec; extension at 72 °C for 30 sec (30 cycles); with final extension at 72 °C for 5 min (1 cycle). The fragment was cloned upstream of the SV40 promoter into the pGL3 luciferase reporter vector (Promega) digested with KpnI-HF and NheI-HF restriction enzymes (New England Biolabs) using NEBuilder^®^ HiFi DNA Assembly (New England Biolabs). The assembled DNA was then transformed in NEB^®^ Stable Competent *E. coli* (New England Biolabs) using ampicillin for selection (Sigma-Aldrich) at 100 μg/mL added to LB Broth (Carl Roth) at 30 ℃ overnight. The plasmid was then sent for Sanger sequencing (Microsynth, Germany) to confirm the correct cloning fragment. To generate a variant (rs77340917, G>A allele) in this plasmid, we employed site-specific mutagenesis method, where the GGAA core motif was changed to AGAA. The primers used are listed in **Supplementary Table S5**. To assess the potential enhancer activity of these constructs in dependency of the *TMPRSS2::ERG* fusion, VCaP cells (8×10^5^ cells per well) were seeded in duplicate wells in 12-well plates and transfected with siPOOLs targeting *ERG* or control siPOOLs. At 48 h post-transfection with siPOOLs, the cells were transfected with the pGL3 reporter assay along with *Renilla* (100:1) using Lipofectamine LTX Reagent with PLUS Reagent (Invitrogen). After a further 48 h, cells were lysed with the 1× Promega lysis buffer and *Firefly* and *Renilla* luciferase activities was recorded using GloMax Discover System plate reader (Promega).

### Statistics

Statistical analysis of both *in vitro* and *in silico* data was performed using GraphPad PRISM 9. Data were presented as mean ± standard error of the mean (SEM). For functional *in vitro* experiments, a two-tailed Mann-Whitney test or a t-test was utilized if not otherwise specified in the corresponding figure legend. Where applicable, the Bonferroni method was used for correction of multiple testing (*P<*0.05). A *P* value <0.05 was considered as statistically significant.

### Data availability

Original RNA-seq transcriptome profiling data have been deposited at the Gene Expression Omnibus (GEO) under the accession code GSE297793. The mass spectrometry raw files, and the search output tables have been deposited to the ProteomeXchange Consortium *via* the PRIDE partner repository under the following identifier: PXD066564. Reviewer access details to PRIDE: Username, Password: HV8QQkjQY1oa.

## LIST OF ABBREVIATIONS

CV: crystal violet
CDF: chip description files
cDNA: complementary DNA
DEGs: differentially expressed genes
DOX: doxycycline
FC: fold change
GS: Gleason score
GWAS: genome-wide association study
HRP: horseradish peroxidase
NES: normalized enrichment score
PCa: prostate cancer
PI: propidium iodide
PTPRK: receptor-type protein tyrosine phosphatase kappa
qRT-PCR: quantitative real-time polymerase chain reaction
RMA: robust multi-array analysis
SNP: single nucleotide polymorphism
STR: short tandem repeat
TE: TMPRSS2::ERG

## SUPPLEMENTARY FIGURES

**Supplementary Fig. 1.**
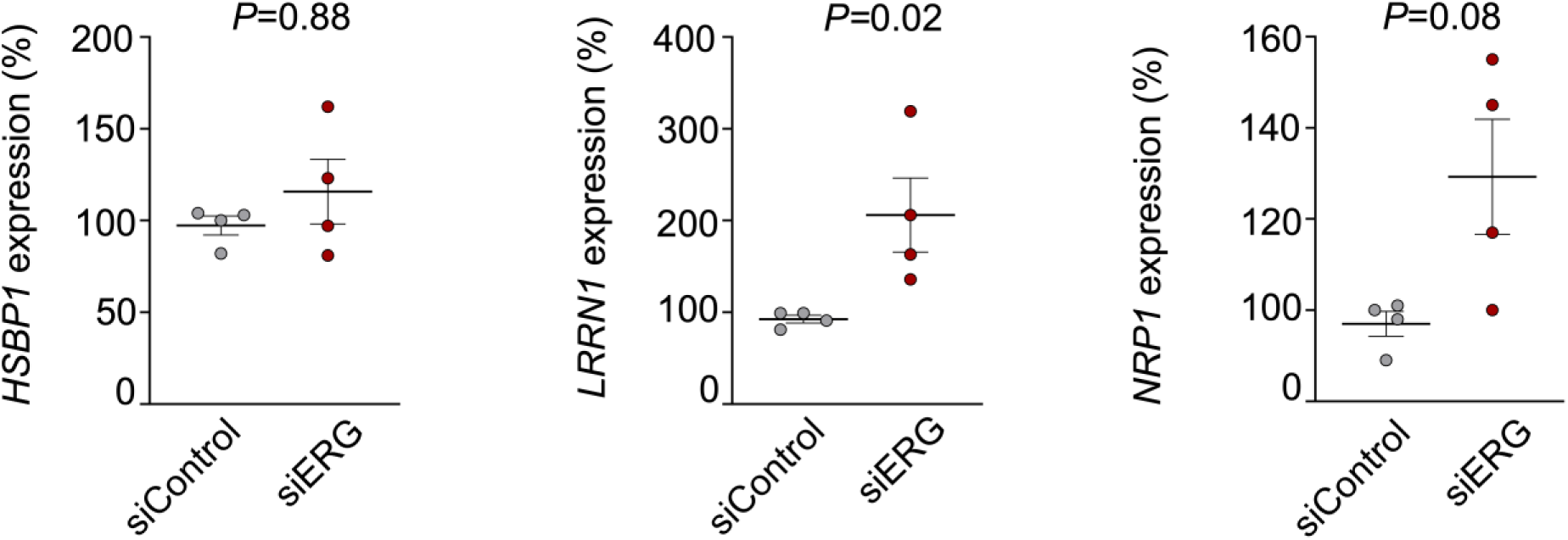
*ERG* knockdown in VCaP cell line and its effect on the expression of *HSBP1*, *LRRN1* and *NRP1*. *HSBP1*, *LRRN1* and *NRP1* expression shown as a percentage of siControl in VCaP cells quantified by qRT-PCR 72 h following siRNA mediated knockdown of *ERG*. Horizontal bars represent means, whiskers represent SEM, n=4 biologically independent experiments. Two-sided Mann-Whitney test.

**Supplementary Fig. 2.**
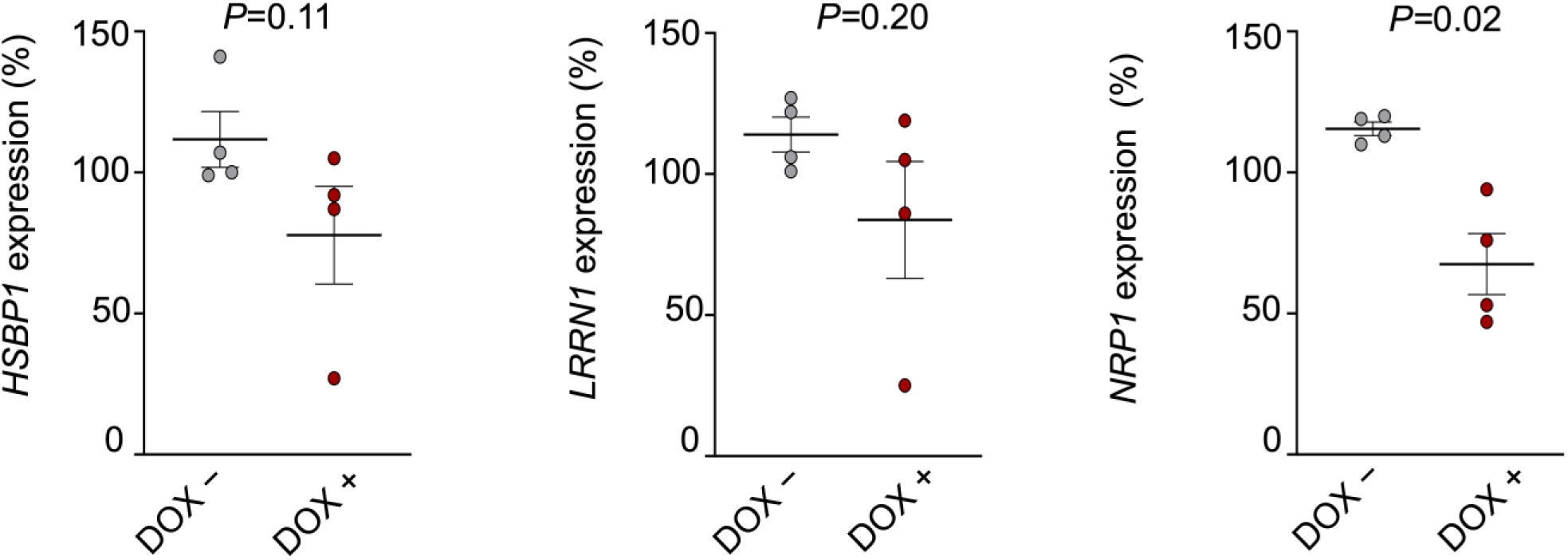
*ERG* overexpression in RWPE-1 cells and its effect on the expression of *HSBP1, LRRN1* and *NRP1*. *HSBP1*, *LRRN1* and *NRP1* expression shown as a percentage of siControl in RWPE-1 prostate epithelial cells expressing a DOX-inducible mediated overexpression of *ERG* at 72 h treatment with/without 1 μg/mL DOX, n=4 biologically independent experiments. Horizontal bars represent means, whiskers represent SEM. Two-sided Mann-Whitney test.

**Supplementary Fig. 3.**
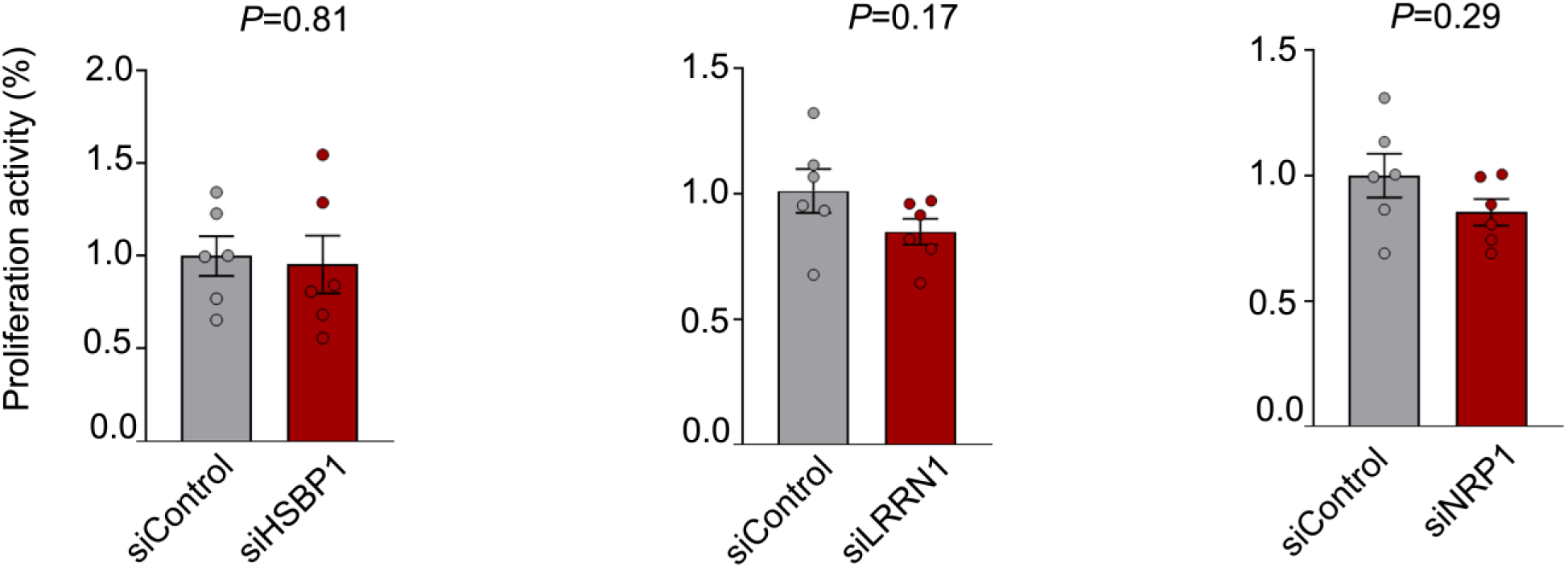
Silencing of *HSBP1, LRRN1 and NRP1* and their effect on proliferation of DuCaP PCa cells. Viable cells count 72 h after transfection shown as a percentage of siControl in DuCaP PCa cell line with siRNA against *HSBP1, LRRN1* and *NRP1* or non-targeting siControl. Horizontal bars represent means, whiskers represent SEM, n=6 biologically independent experiments. Two-sided Mann-Whitney test.

**Supplementary Fig. 4.**
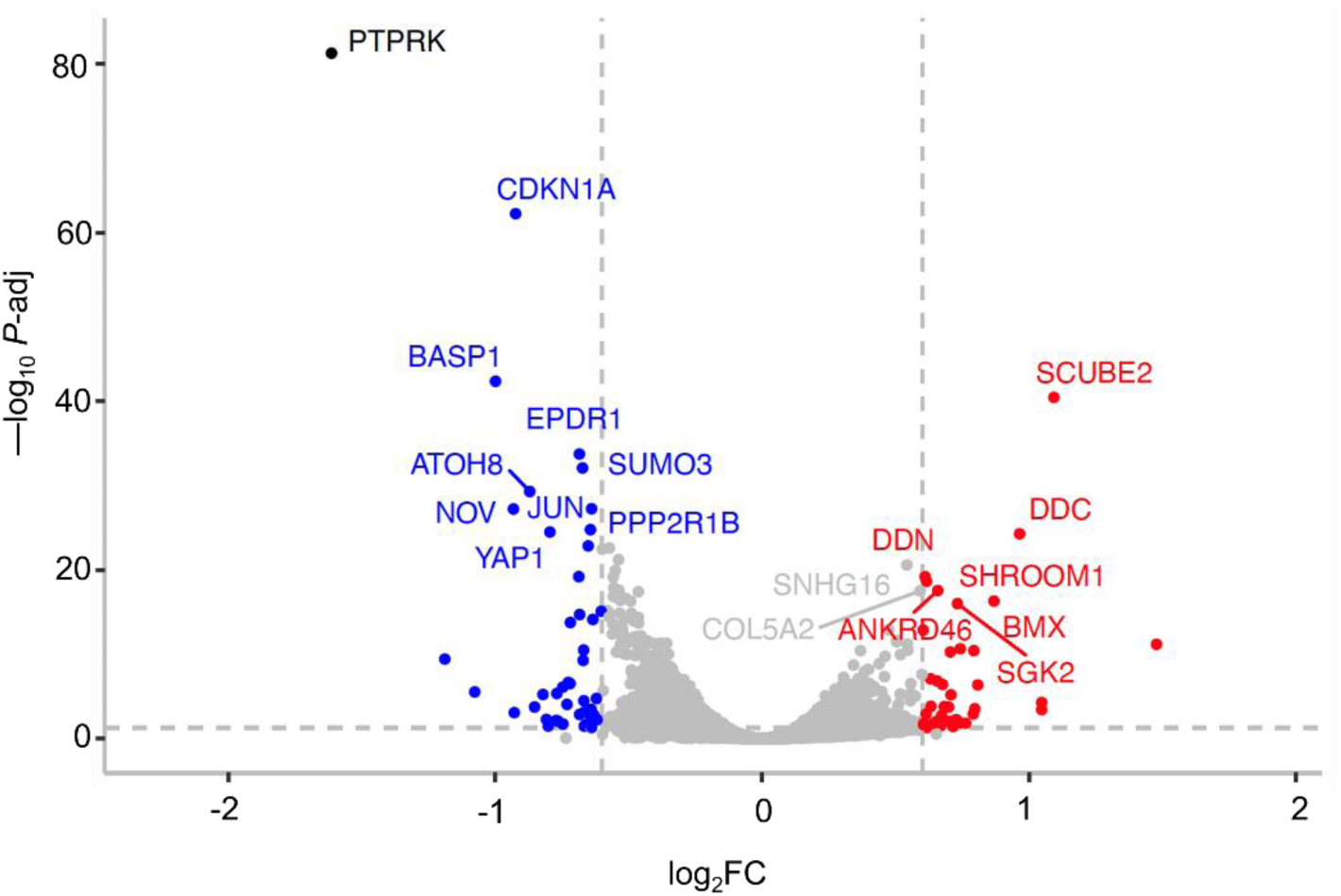
Volcano plot illustrating results of differential gene expression analysis of RNA-seq samples 72 h after transfection of the VCaP cells with siRNA against *PTPRK* or siControl. Blue dots represent down-regulated genes and red dots represent up-regulated genes |log_2_FC| > 0.5 and -log_10_ *P-*adj < 0.01.

**Supplementary Fig. 5.**
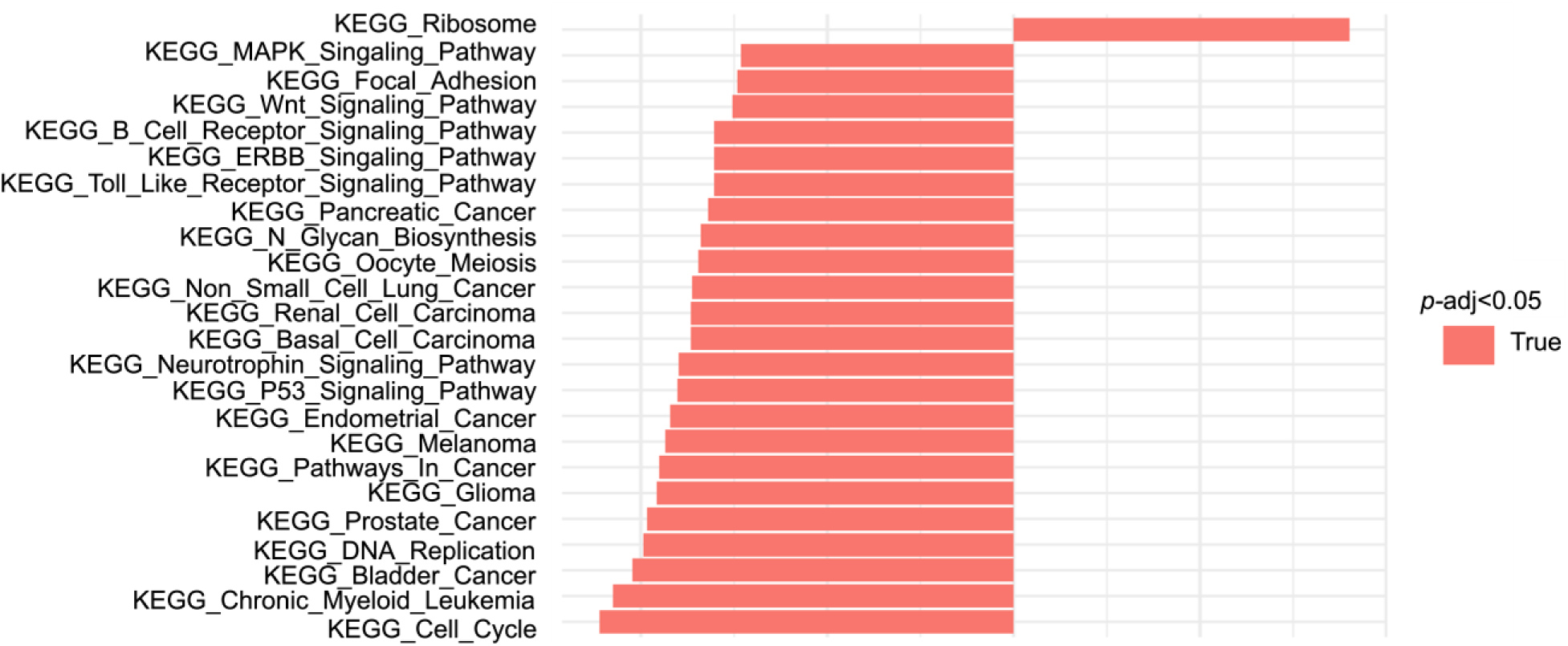
KEGG enrichment analysis of DEGs in VCaP cells 72 h after siRNA mediated silencing of *PTPRK*. Red bars indicate pathways being up/down-regulated upon *PTPRK* silencing.

**Supplementary Fig. 6.**
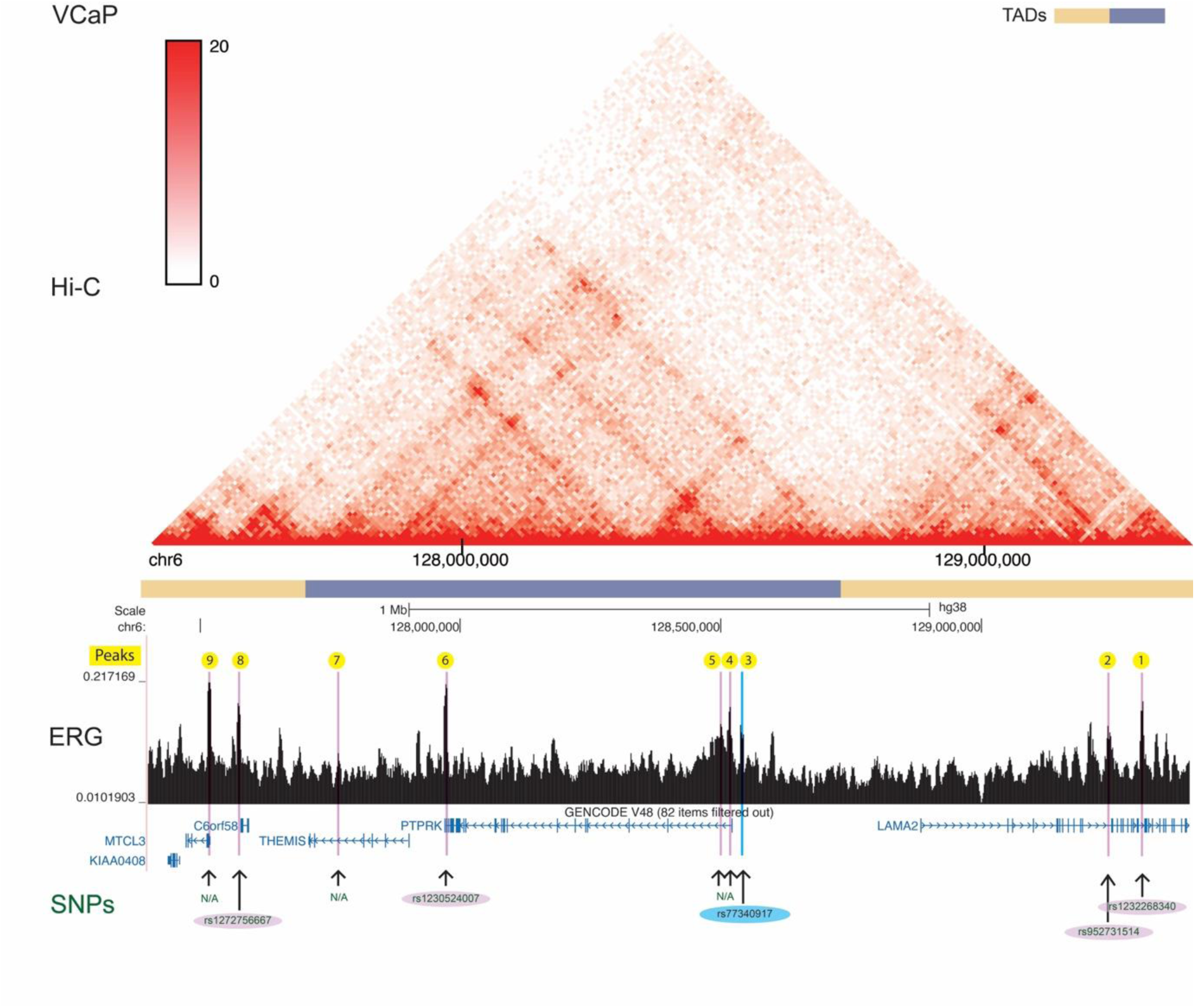
Integrated genomic view of TE ChIP-seq and Hi-C data of the chr6 *PTPRK* locus in VCaP PCa cell model (GSE172097). A total of 9 TE binding sites looping to the *PTPRK* promoter were identified using a publicly available dataset (GSE172097).

**Supplementary Fig. 7.**
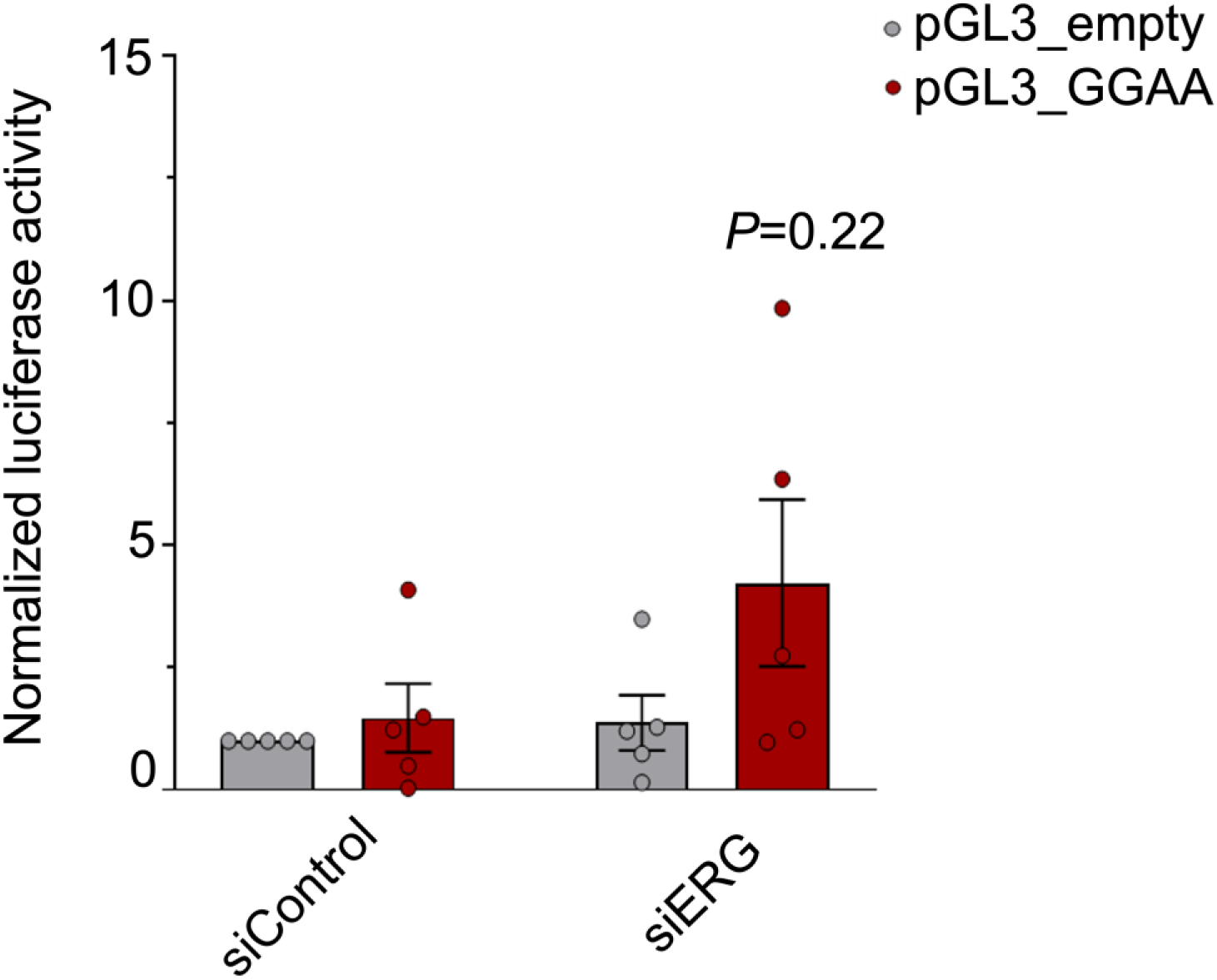
Analysis of the enhancer potential of peak number 2, 72 h after transfection of the VCaP PCa cells with siRNA against *ERG* or non-targeting siControl. Horizontal bars represent mean, whiskers represent SEM, n=5 biologically independent experiments. Two-sided Mann-Whitney test.

**Supplementary Fig. 8.**
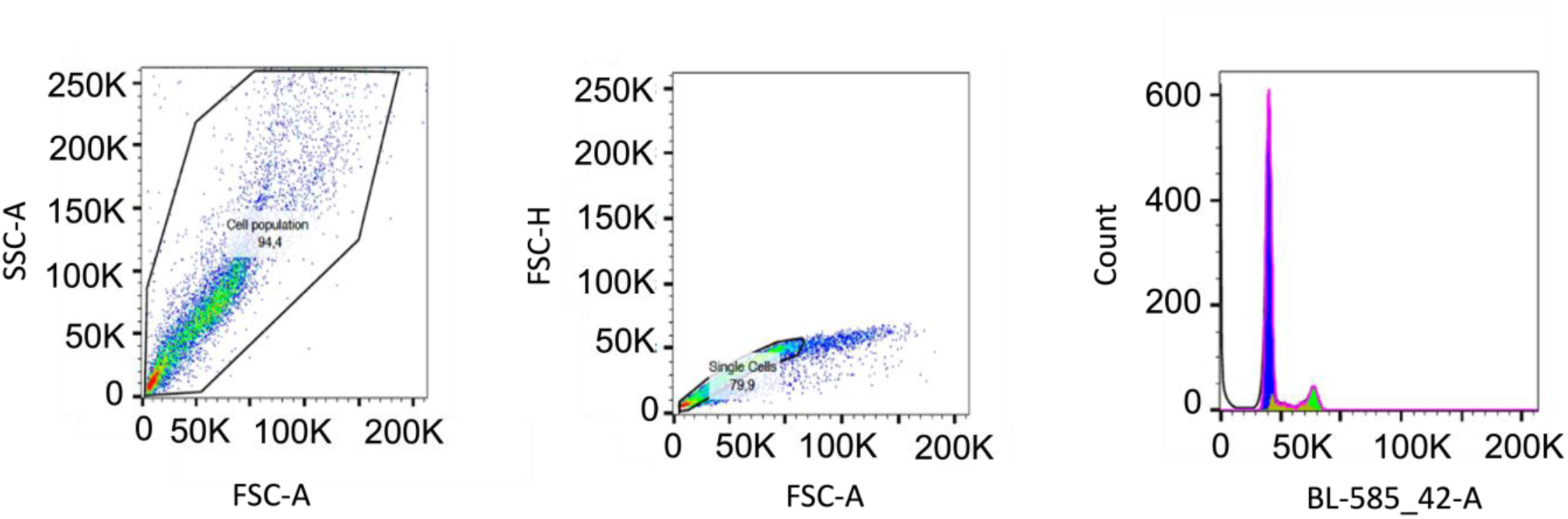
Gating strategy for the analysis of cell cycle (PI, flow cytometry)

